# Delineating plant responses to the 3’,5’- and 2’,3’-cAMP isomers

**DOI:** 10.1101/2025.10.09.681331

**Authors:** Eleonora Davide, Guido Domingo, Angela Di Iacovo, Milena Marsoni, Ping Yun, Chris Gehring, Elena Bossi, Sergey Shabala, Marcella Bracale, Candida Vannini

## Abstract

Similar to animals, both the 3’,5- and the 2,’3’-cAMP isomers are present in plants. The former is the enzymatic product of adenylate cyclases (ACs), the latter is an RNA degradation product. While there is increasing evidence that both isomers can elicit or modulate a broad range of physiological responses, the question of isomer specificity of responses has remained largely unresolved. To delineate isomer-specific responses in *Arabidopsis thaliana* at the systems level, we have combined a comparative proteomics and electrophysiological approaches. Both isomers cause distinct systemic effects on the proteome, with the 2’,3’ isomer notably affecting systems-level functions like transcriptional regulation. None of the isomers affects net ion fluxes in the root under control conditions, but both were able to attenuate the magnitude of oxidative stress-induced K^+^ net loss and Ca^2+^ uptake by 2-fold. Isomer-specific responses of single molecular targets were assessed in the cyclic nucleotide-gated channels 2 and 18 (CNGC2 and CNGC18). Both channels are gated by the 3’,5’-cAMP isomer only, suggesting that the gating is isomer-specific and this implies that gating *in vivo* depends on catalytically active ACs.

**Highlights:** Distinct Arabidopsis responses to 3’,5’- and 2’,3’-cAMP uncover isomer-specific molecular targets and physiological effects in cyclic nucleotide signaling.

## Introduction

Cyclic nucleotide monophosphates (cNMPs) are increasingly recognised as key signalling molecules that directly or indirectly elicit or modify many diverse physiological processes in plants. Among these cNMPs, the cyclic adenosine monophosphate (cAMP) comes in two isomers, 3’,5’-cAMP and 2’,3’-cAMP. The first, 3’,5’-cAMP, is the catalytic product of adenylate cyclases (ACs) that convert ATP into cAMP and pyrophosphate. The second one is the positional isomer 2’,3’-cAMP and is an RNA degradation product catalysed by ribonucleases.

Past reports of plant responses to exogenously applied cAMP largely ignored the specificity of the positional isomer used. Hence, uncertainty prevails in reports that describe cAMP-dependent calcium signaling (Kurosaki and Nishi, 1993; Jin and Wu, 1999), potassium channel activation (Li *et al*., 1994), self-incompatibility responses (Tezuka *et al*., 2007), sodium uptake (Maathuis and Sanders, 2001) and the roles in phytoalexin accumulation (Zhao *et al*., 2004), phenylpropanoid pathway regulation (Pietrowska-Borek and Nuc, 2013), and root heat tolerance (Zhao *et al*., 2021). Recent exclusive studies of 2’,3’-cAMP in plants recognised the 2’,3’ isomer of cAMP as a stress-associated signalling molecule. Functional studies have shown that 2’,3’-cAMP accumulates in response to wounding, heat stress, darkness, and cold (Van Damme *et al*., 2014; Kosmacz *et al*., 2018; Luo *et al*., 2024). In *A. thaliana*, 2’,3’-cAMP treatment can trigger extensive changes in the transcriptome, proteome, and metabolome, notably affecting pathways linked to stress responses (Chodasiewicz *et al*., 2022). More recently, it has been demonstrated in rice that exogenous application of 2’,3’-cAMP induces *DREB1C* expression and significantly enhances chilling tolerance. Incidentally, this effect was not observed in response to 3’,5’-cAMP (Luo *et al*., 2024). However, to the best of our knowledge, there are currently no reports of effects elicited by 2’,3’-cAMP on the ion transport and, in particular, on the gating of cyclic nucleotide-gated channels (CNGCs). Here, we set out to compare the effects of 3′,5′-cAMP and 2′,3′-cAMP in *A. thaliana*, with the aim of assigning specific response signatures elicited by the two isomers. We performed a quantitative proteomics analysis on seedlings treated with each isomer to identify isomer-specific changes in protein abundance and to investigate the potential biological roles of different isomers. Additionally, we employed electrophysiological approaches to evaluate the effects of both isomers on net ion transport and cyclic nucleotide-gated channels 2 and 18 (CNGC2 and CNGC18) (Zelman *et al*., 2012). Our findings provide novel insights into common and distinct molecular and physiological roles of the two cAMP isomers in plant signalling.

## Material and Methods

### Plant material

*A. thaliana* (Columbia ecotype) seeds were surface-sterilized using 70% ethanol (v/v) and 60% bleach (v/v) and then washed five times in sterile water. After two days of stratification in darkness at 4□°C, seeds were inoculated into liquid Murashige and Skoog (MS) medium supplemented with 1% (w/v) sucrose and maintained under continuous light conditions. The medium was replaced after 7 days, and three days later, when seedlings were 10 days old, they were treated with 0.2□µM of the cell permeant Br-2’,3’-cAMP or 0.2□µM Br-3’,5’-cAMP (Chodasiewicz *et al*., 2022). Seedlings treated with water served as control. After 3 hours of treatment, the seedlings were harvested, washed with distilled water for 5 minutes, dried on paper, and flash-frozen in liquid nitrogen; they were then stored at −80°C.

### Workflow of the proteomics analysis

The frozen plant material was ground to a fine powder with liquid nitrogen, and proteins were extracted using the SDS/phenol method (Vannini *et al*., 2021) and digested with trypsin via Filter Aided Sample Preparation (FASP) (Wiśniewski, 2019). Peptides were analysed by LC-MS/MS as detailed elsewhere (Paradiso *et al*., 2020). MS/MS spectra were searched against the *A. thaliana* proteome database (TAIR10, containing 48,266 entries; http://www.arabidopsis.org) using MaxQuant software (version 1.6.10.43) with default parameters (https://www.maxquant.org). For the quantitative proteome, the ‘ProteinGroups’ output file generated by the MaxQuant search was loaded into Perseus software (version 1.6.15.0).

Control versus (vs) 2’3’-cAMP and control vs 3’5’-cAMP treated samples were compared. The two original complete data sets, containing log2-transformed intensities, were split into three subsets each, based on the percentage of replicates with missing or valid values per treatment with a threshold of 75% as described in Supplementary Protocol (Supplementary file 1 and Fig. S1).

### Analyses and data visualization

To generate networks for known and predicted protein–protein interactions, the data sets were loaded to the STRING database (https://string-db.org; version 10.5) using a high confidence interaction score (>0.7). As active interaction sources, text mining, experiments, databases, co-expression, neighbourhood, gene fusion, and co-occurrence were selected. Proteins that did not interact with any other proteins were removed from the network. The network was imported in Cytoscape (version 3.10.3) for protein visualization: we created a clustered version of the network with MCL, and for each cluster, we retrieved enriched functional annotation using the functional enrichment String tool embedded in Cytoscape with default parameters. Only networks with more than three interactions were retained. GO enrichment analysis was conducted using ShinyGO V0.82 (https://bioinformatics.sdstate.edu/go) with the entire species as the background and a p-value cut-off of 0.05. Venn diagrams were created using the Venny 2.1 online tool (http://bioinfogp.cnb.csic.es/tools/venny). To identify potential regulatory mechanisms, transcription factor binding sites (TFBS) within the 1000 bp upstream regions of genes encoding DAPs were analyzed using the web server (http://acrab.cnb.csic.es/TDTHub/; accessed July 1, 2025) TFBS-Discovery Tool Hub (TDTHub) (Grau and Franco-Zorrilla, 2022). The analysis used *A. thaliana* as the reference species and the FIMO algorithm for TFBS mapping, with a minimum S-Score threshold set at 5%.

### Non-invasive Microelectrode Ion Flux Estimation (MIFE) measurement

The net ion fluxes were measured using the non-invasive MIFE technique (University of Tasmania, Hobart, Australia). Specific details on electrode fabrication, calibration, and sample preparation are given elsewhere (Ordoñez *et al*., 2013). Roots of ten-day-old seedlings were immobilized on a cover slide with parafilm strips and immersed in a Petri dish filled with basic salt medium (BSM) solution (0.5 mM KCl and 0.1 mM CaCl2, pH 5.7). K^+^ and Ca^2+^ net fluxes were measured from the root mature zone for 5 minutes as the initial steady state before the application of cAMP. Then, the K^+^ and Ca^2+^ fluxes in response to 10 µM of 8-Br-3’,5’-cAMP or 8-Br-2’,3’-cAMP were monitored for an additional 10 minutes. To investigate the effect of exogenous cAMP on oxidative stress responses, K^+^ and Ca^2+^ net fluxes in response to transient 10 mM H_2_O_2_ were measured from roots of seedlings that were pretreated with or without 8-Br-3’,5’-cAMP or 8-Br-2’,3’-cAMP.

### Preparation of CNGC2 and CNGC18 encoding mRNA

CNGC2 (At5g15410) and CNGC18 (At5g14870) cDNA were synthesised by VectorBuilder and inserted into the pT7 vector within the flanking region of the globin gene of *X. laevis*. The cDNA was amplified in *E. coli* and purified with the Wizard® Plus SV Minipreps DNA Purification System. CNGC2 and CNGC18 cDNAs were linearized by AscI or SgsI, respectively. Corresponding cRNAs were capped and transcribed *in vitro* and using T7 RNA polymerase (Bhatt *et al*., 2022).

### Heterologous protein expression and two-electrode voltage clamp (TEVC) electrophysiology

Electrophysiological recordings were performed in *X. laevis* oocytes purchased from Ecoocyte Bioscience GmbH (Dortmund, Germany). Healthy and full-grown oocytes were microinjected with 25ng/50nL purified cRNA encoding CNGC2 and CNGC18 with a manual microinjection system (Drummond Scientific Company, Broomall, PA, USA). Oocytes were subsequently incubated at 18°C for 3-5 days in NDE solution (96 mM NaCl, 2 mM KCl, 1 mM MgCl_2_, 5 mM 4-(2-hydroxyethyl)-1-piperazineethanesulfonic acid (HEPES) 2.5 mM pyruvate, 0.05 mg/mL gentamicin sulphate, and 1.8 mM CaCl_2_ at pH 7.6) to allow for heterologous expression of CNGC2 and CNGC18. The electrophysiological measurements were performed using a two-electrode voltage clamp (TEVC) at controlled voltage conditions with the Oocyte Clamp OC-725C or B amplifier (Warner Instruments, Hamden, CT, USA). The digitally recorded currents were collected using the pCLAMP software (Version 11.2.2, Molecular Devices, San Jose, CA, USA). The two microelectrodes were filled with 3 M KCl, and bath electrodes were connected to the oocyte chamber via two agar bridges (3% agar in 3 M KCl). To measure K^+^-currents, the external control solution contained 96 mM KCl, 1.8 mM CaCl_2_, 1.8 mM MgCl_2_, and 10 mM HEPES-KOH, pH 7.5. The Ca^2+^-currents were measured using an external control solution composed of 10 mM CaCl_2_, 185 mM D-mannitol, and 10 mM MES-Tris, pH 7.4. The current-voltage (I-V) relationship was obtained using a voltage jump protocol consisting of 10 square pulses (0.8 s long) ranging from −180 mV to 0 mV (in 20 mV increments) before and after incubation (30 minutes) with 50μM of either 8-Br-3’,5’-cAMP or 8-Br-2’,3’-cAMP

### Statistical analysis

For proteomic analysis the statistical workflow was performed as described previously (Nikonorova *et al*., 2018): In brief, only the subset 1, obtained as described in Supplementary Protocol (Supplementary file 1), was centred around zero by subtracting the median within each replicate (n = 4) and subjected to a two-sample t-test (P < 0.05).

For MIFE technique, the statistical significance in H_2_O_2_-induced ion fluxes was determined by One-way ANOVA, Šídák’s multiple comparisons (*p* < 0.05).

For electrophysiology analysis the data were collected using Clampfit 11.2.2 software (Molecular Devices, Sunnyvale, CA, USA). OriginPro 8.0 (OriginLab Corp., Northampton, MA, USA). OriginPro 8 and GraphPad Prism 8.0.2 (GraphPad Software, Boston, MA, USA) were used for statistical analyses. The statistical significance of the data was determined using the unpaired *t*-test. Levels of significance were defined as *, p□< 0.05; **, p□<□0.01; ***, p□< 0.001; ****, p <□0.0001. The number of oocytes (n) obtained from a frog (N) used in each experiment is indicated as “n/N”.

## Results

### Early changes in the proteome in response to cAMP isomers

To gain insight into early systemic changes elicited by the two cAMP isomers, we focused on changes in the proteome of *A. thaliana* seedlings after three hours of exposure to cell-permeant derivatives of 3’,5’-cAMP or 2’,3’-cAMP. The Br derivatives of both 3′,5′-cAMP and 2′,3′-cAMP have been successfully applied in studies of cAMP signaling in plants (Jiang *et al*., 2005; Alqurashi *et al*., 2016; Kosmacz *et al*., 2018; Chodasiewicz *et al*., 2022)

We selected a concentration of 200 nM for both isomers, corresponding to the same order of magnitude as the intracellular level reported for 3′,5′-cAMP (Ashton and Polya, 1978; Moutinho *et al*., 2001).

Proteomic analysis of control and cAMP-treated samples identified 5614 protein groups in total (Tab. S1). We applied a hybrid analytical approach that separately considers intensity-based and presence/absence data (Nikonorova *et al*., 2018) and categorized the dataset into three subsets (Fig. S1, Supplementary file 1). In subset 1, we included proteins with sufficient quantitative data for statistical analysis. Subset 2 contained proteins with unreliable or inconsistent detection levels, and subset 3 contained “unique” proteins that were present under one condition only (either treatment or control). Furthermore, we classified the differentially abundant proteins (DAPs, Tab. S2 and S3) in up-regulated (detected only or significantly more abundant in cAMP-treated samples) and down-regulated proteins (detected only in control samples or significantly more abundant under control conditions). In a total of 168 DAPs in 3’,5’-cAMP-treated plantlets, 97 were upregulated and 71 were downregulated. In 168 DAPs identified from the 2’,3’-cAMP-treated plantlets, 49 were upregulated and 119 were downregulated. A Venn diagram revealed that only 33 DAPs are shared between the two treatments (Fig. 1A, Tab. S4). The majority of common proteins are annotated as having roles in metabolic processes, including protein metabolism and response to stimuli (Fig. 1B, Tab. S4).

**Fig. 1.**
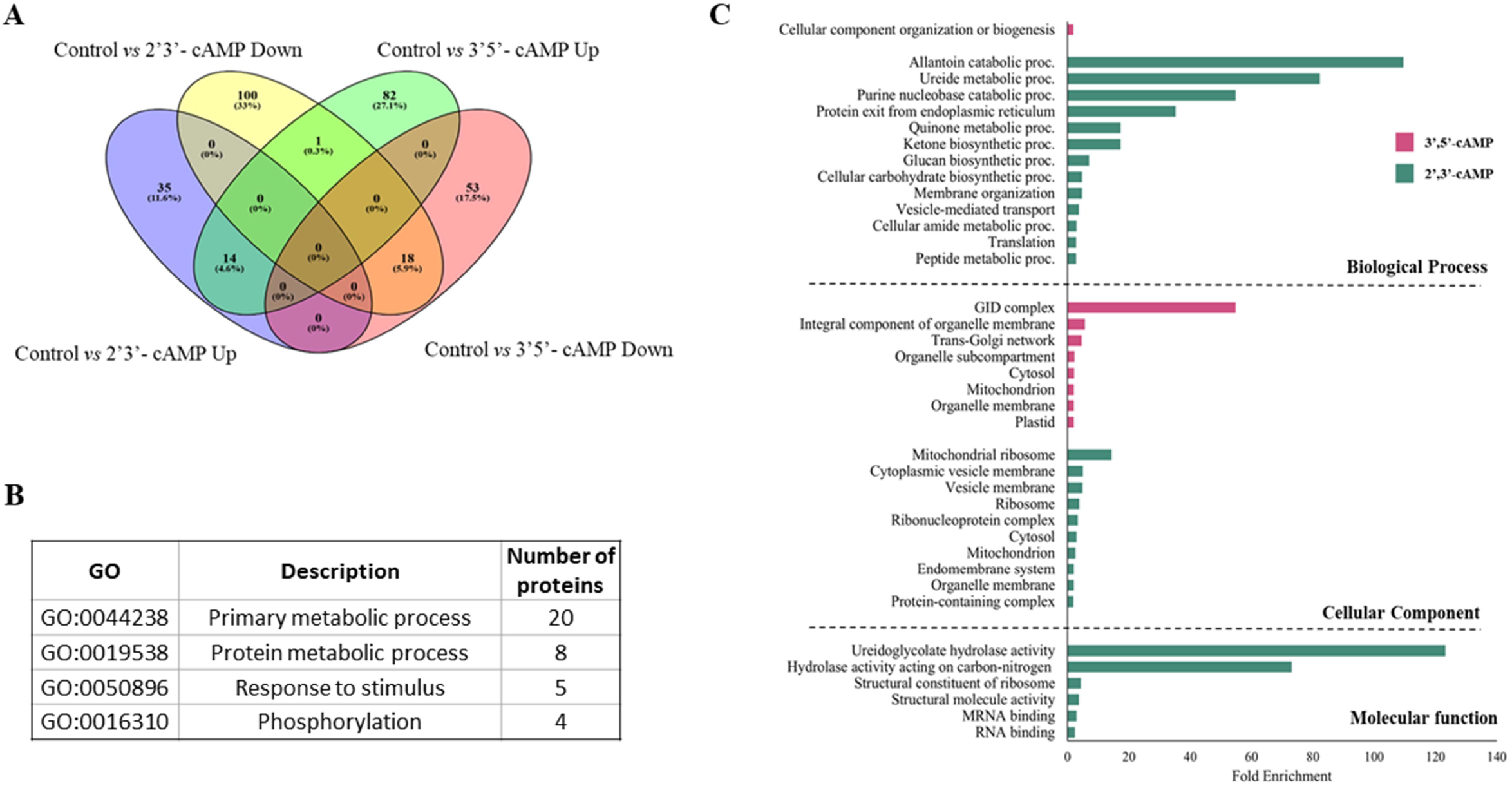
A comparative proteomics after 2′,3′-cAMP and 3′,5′-cAMP treatment. **A**. Venn diagram showing the overlap of significantly regulated proteins following 0,2 µM Br-2′,3′-cAMP or Br-3′,5′-cAMP treatment. **B**. Functional classification of the shared DAPs. **C**. Gene Ontology term enrichments of biological process, cellular Additional, and molecular function were significantly overrepresented among the differentially abundant proteins (FDR-corrected p ≤ 0.05, n = 4 biological replicates).

Gene Ontology (GO) enrichment analysis of the DAPs was conducted using ShinyGO V0.82 (based on Ensembl Release 104, Feb. 3, 2025) (Ge *et al*., 2020) (Fig. 1C). Biological process analysis showed that DAPs linked to 3’,5’-cAMP treatment were enriched in cellular component organization or biogenesis. In terms of cellular component enrichment, these DAPs were associated with the Golgi apparatus, cytosol, mitochondrion, and plastid. No significant enrichment was observed for molecular functions. In turn, DAPs from the 2’,3’-cAMP treatment were significantly enriched in biological processes such as purine nucleobase catabolic process, quinone metabolic process, vesicle-mediated transport, and translation. (Fig. 1C). Cellular components analysis revealed enrichment in the ribosome, vesicle membrane, cytosol, and mitochondrion. Molecular function enrichment indicated that these DAPs were involved in ureide hydrolase activity, structural components of the ribosome, and mRNA binding.

To gain further insight into the relationships among the DAPs in response to the isomers, we constructed a protein-protein interaction (PPI) network using String with a high confidence interaction score. This analysis revealed that the majority of interacting DAPs in both treatments were associated with translation, RNA binding, and ribosomal activity (Fig. 2). Notably, 3′,5′-cAMP treatment led to an increased abundance of the proteins related to these processes, whereas 2′,3′-cAMP treatment decreased the abundance of proteins involved in these functions. Additional cluster associated with ribosome biogenesis was specifically linked to 3’,5’-cAMP (Fig. 2A), whereas the 2,3’-cAMP-associated DAPs formed clusters related to splicing and ribonucleoprotein complexes, oxidative phosphorylation, and vesicle-mediated transport (Fig. 2B).

**Fig. 2.**
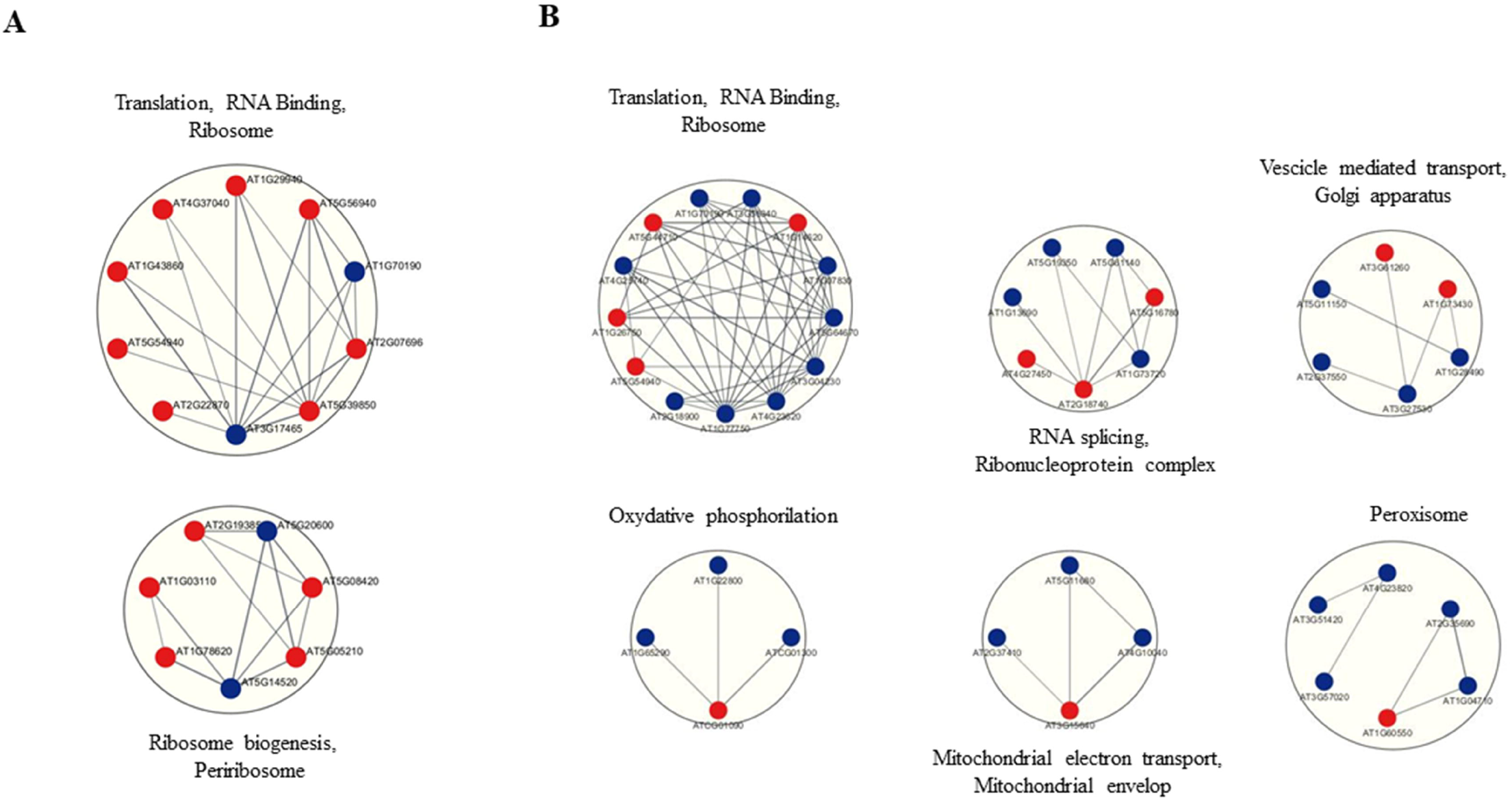
Functional protein-protein interaction networks. **A**. DAPs induced by 3’5’-cAMP **B**. DAPs induced by 2’3’-cAMP. DAPs upregulated were indicated with red circles, while the downregulated were shown with blue circles.

To infer common and specific cAMP-isomer-dependent mechanisms, transcription factor-binding sites in the upstream regions of genes encoding differentially accumulated proteins were analyzed using the TFBS-Discovery Tool Hub (TDTHub) web server (Grau and Franco-Zorrilla, 2022). Upstream regions of genes encoding unique 3’,5’-cAMP-upregulated DAPs appear significantly enriched in consensus *cis*-elements for transcription factors, including CMTA3, DOF5.4, and GBF1. In contrast, no significant enrichment was observed in the upstream regions of genes encoding the 2’,3’-cAMP-upregulated DAPs (Tab. S5).

### 3’,5’- and 2’,3’-cAMP modulate oxidative stress-induced K□ and Ca^2^□ net fluxes in *A. thaliana* roots

To explore the physiological roles of the two cAMP isomers on Arabidopsis roots, the MIFE technology for non-invasive ion flux measurements was applied to monitor real-time net fluxes of K□ and Ca^2^□ from the mature root zone with high temporal resolution (5 seconds). Seedlings were preincubated for 30 minutes with either isomer or with a control (BMS) solution under non-stress conditions. No difference in basal levels of K□ or Ca^2^□ fluxes between controls and roots treated with either 3’,5’-cAMP or 2’,3’-cAMP was found (Fig. 3 inset). Roots’ exposure to oxidative stress resulted in a massive net K+ loss and Ca2+ uptake (Fig. 3). Root pretreatment with either 3’,5’-cAMP or 2’,3’-cAMP significantly attenuated H□O□-induced ion fluxes, reducing the response magnitude to approximately 50% of that observed in untreated (mock control) roots (Fig. 3). These findings demonstrate that both isomers can directly or indirectly modulate net fluxes and mitigate ionic imbalances caused by H_2_O_2_.

**Fig. 3.**
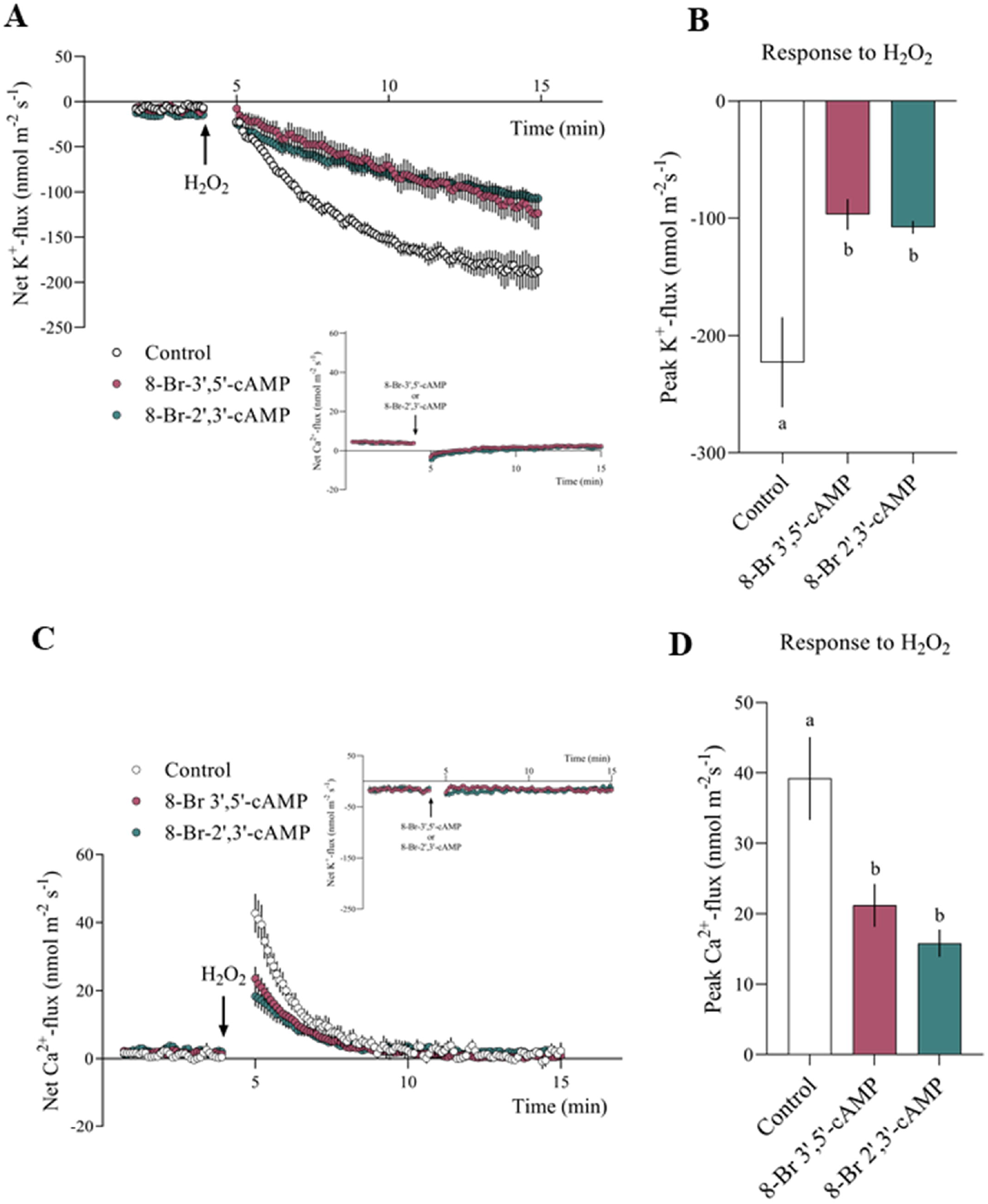
Modulatory effects of 2’,3’- and 3’,5’-cAMP on H_2_O_2_-induced ion fluxes in *A. thaliana* roots. **A**: Transient K^+^ flux; **B**: peak of net K^+^ efflux (nmol m^-2^ s^-1^); **C**: Transient Ca^2+^ flux; **D:** peak of net Ca^2+^ flux (nmol m^-2^ s^-1^). Data are mean ± S.E. (n= 5-9 individual plants). Transient treatment with solely 8-Br-3’,5’-cAMP or 8-Br-2’,3’-cAMP didn’t affect ion fluxes (insets of net K^+^ and Ca^2+^ flux graphs). DiLerent lowercase letters indicate a significant diLerence in H_2_O_2_-induced ion fluxes at *p* < 0.05 (One-way ANOVA, Šídák’s multiple comparisons test). The sign convention of ion flux is “negative efflux”.

### Testing the cAMP isomers as modulators of CNGC2 and CNGC18

CNGC2 and CNGC18 have been reported to be gated by 3’, 5’-cAMP when expressed in *X. laevis* oocytes (Leng *et al*., 1999). This is likely due to the presence of a conserved region, the phosphate binding cassette (PBC), which interacts with the sugar and phosphate moieties of cNMP ligands. A “hinge” region between the PBC and the CaMBD may contribute to ligand binding efficacy and selectivity (Zelman *et al*., 2012). Since the PBC binds exclusively to the sugar and the phosphate group of cNMPs, it may represent a specific target for only one isomer. Here, we employed an electrophysiological approach to determine if the response to cAMP in these two CNGCs is isomer-specific.

To test this, RNA encoding for CNGC2 or CNGC18 channel was injected into Xenopus oocytes, and whole-cell current recorded by TEVC using a voltage step protocol from the holding potential of −60 mV (20 mV step, from –180 mV to 0 mV) (Fig. 4). Since acute perfusion with 3′,5′-cAMP in the recording chamber, was insufficient to elicit any currents, we employed a pre-incubation protocol (Leng *et al*., 1999) in which oocytes expressing the channel were incubated for 30 minutes with either 50 µM of 3′,5′-cAMP or 50 µM 2′,3′-cAMP. Currents of CNGC2 and CNGC18 expressing oocytes were recorded before and after the incubation. In the oocytes expressing both CNGC2 and CNGC18, an inward cationic current was observed following incubation with 3′,5′-cAMP, both in the presence of either K^+^ (96 mM KCl; Fig. 4A-4C-red) or Ca^2+^ (10 mM CaCl; Fig. 4B-4D red).

**Fig. 4.**
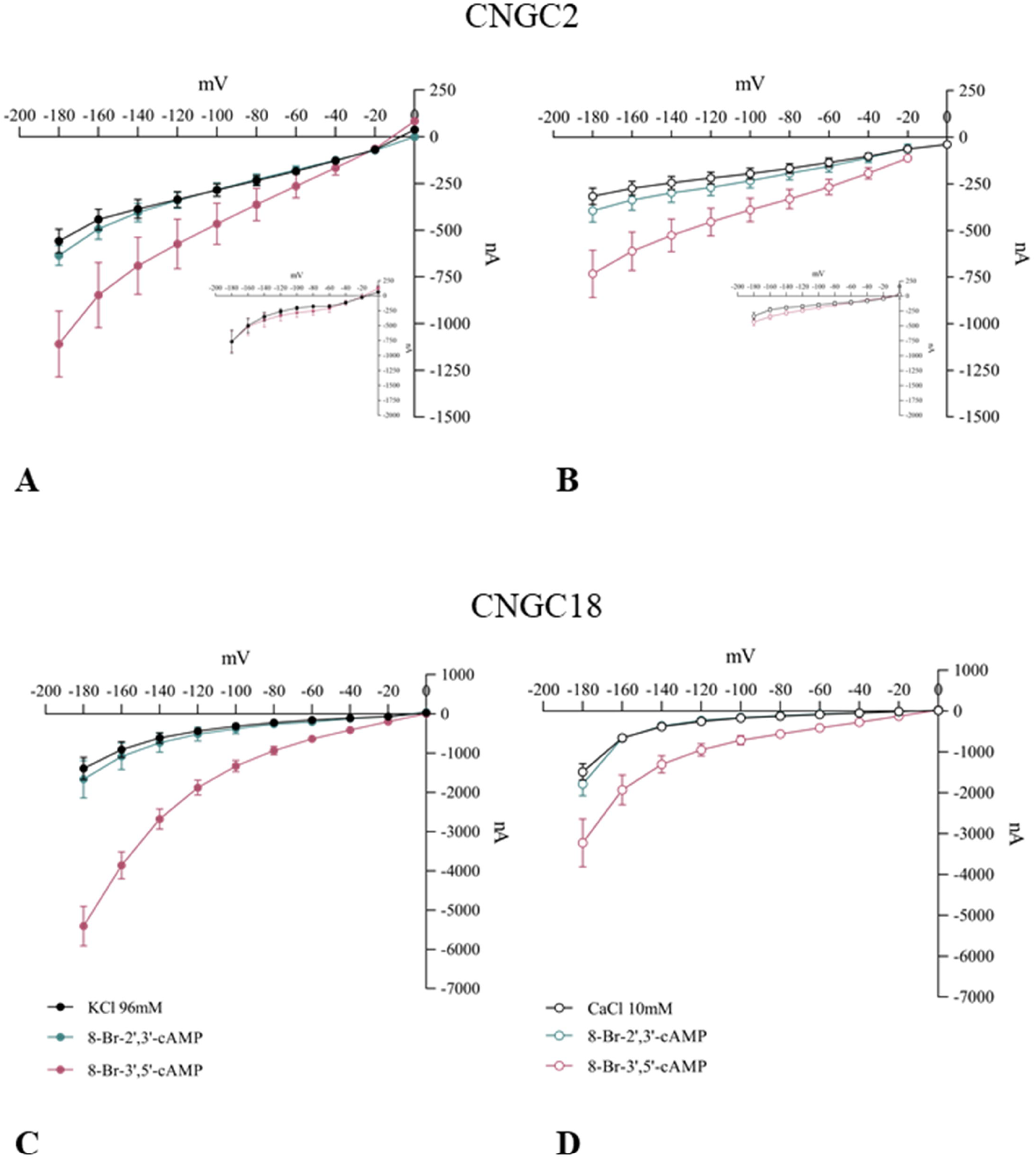
CNGC2 and CNGC18 are activated by 3′5′-cAMP but not by 2′3′-cAMP. **A-B** I-V relationship recorded in oocytes expressing CNGC2 under the same conditions as in A. (n/N=15/2; n/N=11/2) and B. (n/N=13/2; n/N=11/2). Non-injected oocytes exhibit no response to 3′5′-cAMP under either condition (insets of K^+^- and Ca^2+^-current graphs). **C** I-V relationship (from −140 mV to 0 mV) in oocytes expressing CNGC18 shows inward K^+^-currents in response to 3′5′-cAMP (n/N=11/2), while no activation is observed with 2′3′-cAMP (filled dots) (n/N=9/2). **D** As in A., Ca^2+^-currents (open dots) are observed with 3′5′-cAMP (n/N = 8/2) but not with 2′3′-cAMP (n/N = 10/2).

When the ligand was 2′,3′-cAMP, however, no current was observed (Fig. 4). This is clearly visualized by analyzing the net current values obtained by subtracting the baseline current recorded from the current recorded after 30 minutes (Fig. 5). As expected, non-injected oocytes did not show any response after the incubation in 3′,5′-cAMP either in Ca^2+^- or K^+^-containing solutions (Fig. 4 inset). In order to further define the current activation lag phase, we performed a time course by microinjecting 50 nL of 3’,5’-cAMP (50 µM) into oocytes expressing CNGC2 and monitoring the KL current (96 mM KCl) at 10-minute intervals. Under these conditions, we observed maximal channel opening between 20 and 40 minutes (data not shown). Doubling the amount of 3’,5’-cAMP (100 µM) did not shorten the lag phase.

**Fig. 5.**
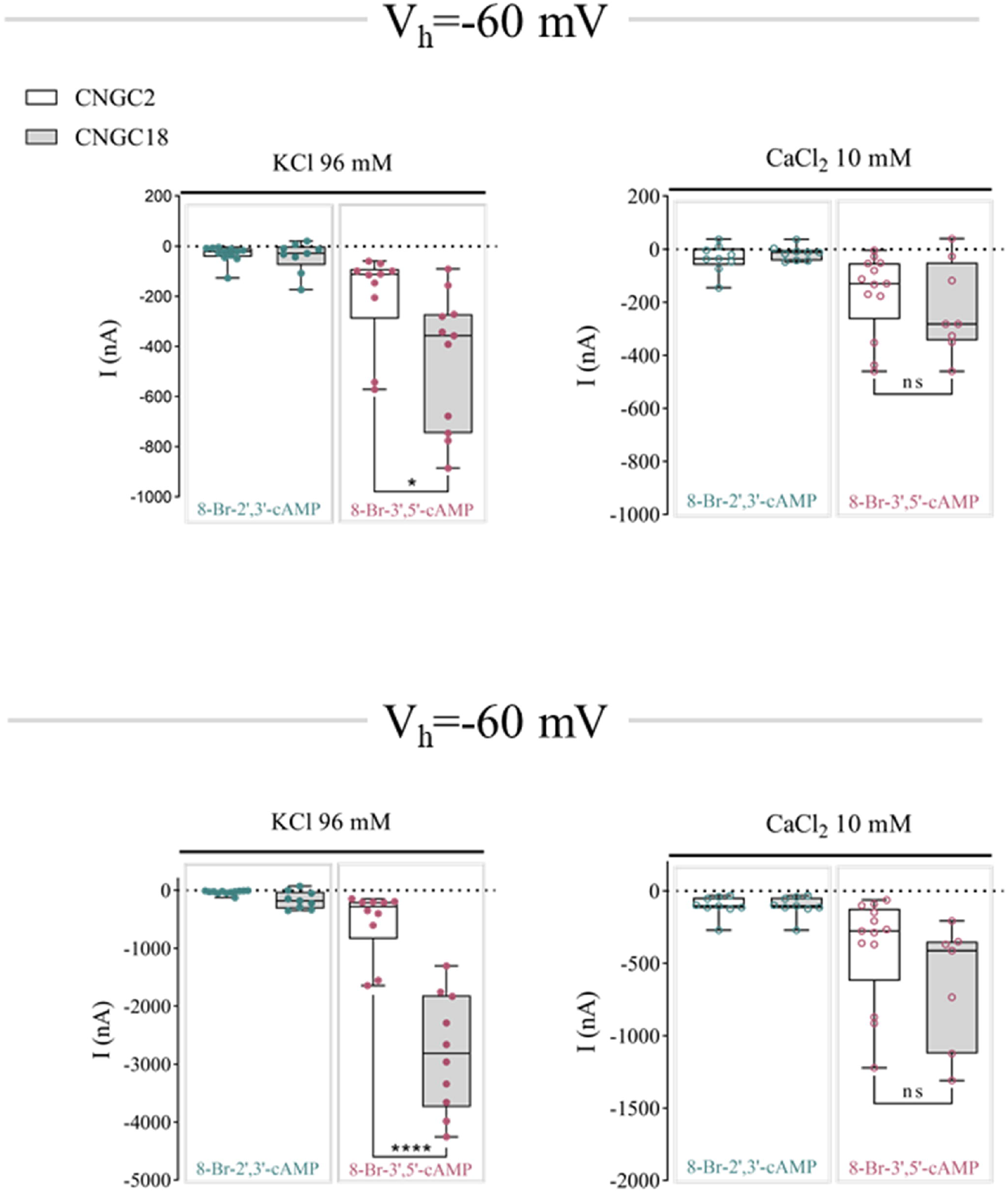
Box-plots representing the current amplitudes elicited in oocytes after 30 min of incubation in 2’3’cAMP and 3’5’cAMP. The net currents were calculated by subtracting the basal current (recorded in the presence of KCl 96mM or CaCl_2_ 10mM) from the substrate-induced current.

## Discussion

The cyclic nucleotide isomers 3′,5′-cAMP and 2′,3′-cAMP differ markedly in their biosynthetic origins. The 3′,5′-cAMP isomer is produced by ACs, which in plants are often embedded within multifunctional proteins (Al-Younis *et al*., 2015, 2018; Chatukuta *et al*., 2018; Bianchet *et al*., 2019; Ruzvidzo *et al*., 2019). Some of these ACs can be activated as part of receptor-linked responses to diverse extracellular cues, positioning 3′,5′-cAMP as a localized and dynamic signal that regulates key physiological processes, including ion homeostasis, stomatal movement, pollen tube growth, and stress adaptation (Gehring and Turek, 2017; Blanco *et al*., 2020). In contrast, 2′,3′-cAMP is generated during RNA degradation and is therefore enhanced under stress conditions, such as wounding, heat, or hypoxia, resulting in increased intracellular levels of 2′,3′-cAMP (Thompson *et al*., 1994; Jackson *et al*., 2009; Van Damme *et al*., 2014). The link between stress-induced RNA turnover and 2′,3′-cAMP accumulation may indicate a role as a cellular effector or response metabolite, rather than as a signalling molecule in the classical sense. 2′,3′-cAMP may provide and quantitative, dose response-like link to stress damage, also raising the possibility that 2’,3’-cAMP functions in cellular protection and metabolic adaptation. The discovery that Toll/Interleukin-1 receptor (TIR) domains can act as 2′,3′-cAMP/cGMP synthetases by hydrolysing RNA/DNA implies that this isomer may be generated in highly regulated signalling contexts, particularly in plant immunity (Yu *et al*., 2022). Furthermore, in plants the presence of both isoform-specific and dual-specificity phosphodiesterases (Lin and Varner, 1972; Brown *et al*., 1977; Tyc *et al*., 1987; Diffley *et al*., 2001) is consistent with the calibration of isomer levels and, consequently, the tuning of their downstream effects.

Our systems-level analysis, conducted using proteomics, has revealed that the two cAMP isomers result in a markedly different set of DAPs, with just 20% of proteins shared. This implies that both isomers largely affect specific biological pathways and GO enrichment and PPI analyses emphasize this difference. Notably, the 3′,5′-isomer predominantly influenced target proteins associated with roles in cellular component organization and biogenesis, process typically linked to the regulation of growth and development. The 2’,3’-isomer affects proteins with a role in purine nucleobase catabolism, which could be interpreted as a feed-back stimulation in response to the nucleotide addition and suggests that stress-induced RNA degradation producing 2’,3’-cAMP will eventually also stimulate nuclease degradation to prevent flooding the system with 2’,3’-cAMP. The isomer effect of quinone metabolism is also noteworthy given their role in cellular respiration, photosynthesis, the metabolism of xenobiotics, and the production of phenylpropanoids.

Importantly, quinones serve as key redox-active compounds. Their dual nature enables them to generate reactive oxygen species (ROS), which can cause oxidative damage, but they also function as antioxidants and signalling molecules, helping plants detect and respond to stress (Lüthje *et al*., 2013; Kozuleva *et al*., 2014).

Changes in the vesicle-mediated transport are also part of systemic stress responses that, in turn, cause elevated cellular 2’,3’-cAMP levels (Wang *et al*., 2020; Sampaio *et al*., 2022, Preprint). The regulation of transcription is also a systems-level response, and again significantly affected by 2’,3’-cAMP treatment. The prominence of RNA-binding proteins in the 2′,3′-cAMP-treated samples is entirely consistent with previous reports linking this isomer to stress granule dynamics and mRNA turnover (Kosmacz *et al*., 2018; Chodasiewicz *et al*., 2022). PPI analysis supports these connections and points to the operation of networks that integrate translation, RNA binding, and ribosomal function. Additional clusters unique to 2′,3′-cAMP included splicing and ribonucleoprotein complex formation, which is again diagnostic for a systemic effect of 2′,3′-cAMP on RNA processing. Finally, the downregulation of proteins involved in oxidative phosphorylation and mitochondrial transport has also been observed in animal cells, where elevated levels of 2′,3′-cAMP have been implicated in mitochondrial dysfunction (Azarashvili *et al*., 2009). Furthermore, transcription factor (TF) binding site analyses again point to distinct mechanisms of action for the two isomers. Upstream regions of genes encoding 3’,5’-cAMP-upregulated DAPs appear significantly enriched in *cis*-elements for transcription factors, including CMTA3, DOF5.4, and GBF1. These *cis*-elements are not enriched in genes encoding 2’,3’-cAMP-upregulated DAPs. Notably, the calmodulin-binding transcription factor CMTA3 had previously been proposed as a TF regulated by 3’,5’-cAMP during the Arabidopsis immune response (Sabetta *et al*., 2019).

To further study the system-wide effects of two cAMP isomers, we quantified the ion net fluxes from roots. Extensive literature exists regarding the impact of 3’,5’-cAMP on ionic flux under stress conditions (Ordoñez *et al*., 2014). However, no such data are available for 2’,3’-cAMP. It is well established that ROS can activate both outward-rectifying potassium channels and non-selective cation channels (NSCCs) in the root epidermis, and this can cause net K□ efflux and Ca^2^□ influx (Demidchik *et al*., 2003, Demidchik *et al*., 2010; Shabala and Pottosin, 2014). ROS-mediated channel activation is essential in early stress responses and can contribute to physiological adaptation. In our study, a pre-treatment with either 2′,3′- or 3′,5′-cAMP significantly reduced H□O□-induced K□ efflux and Ca^2^□ influx, and this is consistent with cAMP-dependent ROS-activated ion transport. Both isomers cause similar changes to the net-flux signature and may do so via common and/or at least partly different pathways.

In order to assess specific and direct effects of the isomers, we turned to the CNGCs. A hallmark of the 3’,5’-cAMP signalling in plants is its capacity to enable ion flux through specific CNGCs (Zelman *et al*., 2012), such as CNGC2 and CNGC18 (Leng *et al*., 2002; Zhou *et al*., 2014), which have a role in various physiological responses, including immune signalling and pollen tube guidance. To date, reports of cAMP effects on CNGCs have considered the 3′,5′-cAMP isomer only. Here, we have been able to elicit inward cationic currents in Xenopus oocytes expressing either CNGC2 or CNGC18 from *A. thaliana* in the presence of the 3′,5′-cAMP. In contrast, 2′,3′-cAMP failed to induce any measurable current, and this provides direct functional evidence for isomer-specific cAMP responses and suggests that AC activation is an essential upstream part of cAMP-dependent CNGC modulation.

Interestingly, we observed a reproducible lag phase, also when the cAMP was injected, before the activation of 3′,5′-cAMP-induced currents. This phenomenon has already been observed in guard cell protoplasts under similar experimental conditions in the whole-cell configuration(Ali *et al*., 2007). The authors have speculated that the long incubation times required for ligand activation could be due to competition for binding sites between cAMP and CaM. In plant CNGCs, these sites are both located in the C-terminal domain, differing from the animal CNGC configurations. The authors propose that a further reason might be that plant cells have comparatively relatively higher levels of cyclic nucleotide phosphodiesterases than animal cells; however, this point is not relevant in our oocyte system, in which we use non-hydrolyzable cAMP as well as a heterologous animal system. It appears more likely that the lag may be caused by the time required for cAMP to diffuse in a crowded cytoplasmic microenvironment and to reach and saturate the binding sites. Furthermore, these findings are also consistent with the idea that cyclic nucleotide signalling in plants operates within spatially restricted cellular microenvironments where cAMP signalling is tightly localized to prevent non-specific activation and to ensure signal fidelity (Gehring and Turek, 2017). In the context of plant cells, membrane nanodomains and localized pools of cyclic nucleotide-producing enzymes likely contribute to this compartmentalization, where, e.g., moonlighting ACs are located close to cAMP-dependent proteins like CNGCs. The Xenopus oocyte system lacks such spatial constraints, which may account for the observed kinetic delay and provide further support for the physiological relevance of localized signalling *in vivo*.

In summary, our data show that both isomers directly or indirectly cause distinct systemic effects on the proteome, with the 2’,3’ isomer affecting systems-level functions such as transcriptional regulation. None of the isomers affects net ion fluxes in the root under control conditions, but both elicit the same response under H_2_O_2_ stress conditions. The mechanisms by which this occurs will need to be elucidated. Finally, the cAMP gating effect on CNGC2 and CNGC18 is 3’,5’-AMP isomer-specific and, hence, dependent *in vivo* on activated adenylate cyclases.

**Supplementary file 1**. Supplementary Protocol.

**Supplementary Figure S1**. Filtering data system.

**Supplementary Table S1-S4**. Total dataset, DAPs and shared DAPs.

**Supplementary Table S5**. Transcription factor-binding sites enrichment analysis.

## Acknowledgements

E.D. is a Ph.D. student of the “Life Sciences and Biotechnology” course at University of Insubria. This work was supported by the University of Insubria grant “Fondo di Ateneo per la Ricerca 2024 and 2025” (to B.M. and C.V.).

Proteomics was performed with the support of the Functional Genomics Center Zurich (FGCZ) of University of Zurich and ETH Zurich.

## Authors’ contributions

ED and ADI: In Vivo Experiments, Data Collection, Original Draft

ED, ADI and PY: Data analysis;

ED, ADI and SS: Validation;

MM and GD: Software, Computational Analysis, Visualization;

EB and SS: Resources;

CV, CG and EB: Conceptualization;

EB: Methodology Writing;

EB, PY, SS, CV and CG: Writing– Review & Editing;

MB and CV: Funding Acquisition;

PY, SS, CV and CG: Supervision.

## Data Availability

The mass spectrometry data have been deposited to the ProteomeXchange Consortium (http://proteomecentral.proteomexchange.org) via the jPOST partner repository (https://repository.jpostdb.org) with the data set identifier PXD066253 for ProteomeXchange and JPST003946 for jPOST

## References

Ali R, Ma W, Lemtiri-Chlieh F, Tsaltas D, Leng Q, von Bodman S, Berkowitz GA. 2007. Death Don’t Have No Mercy and Neither Does Calcium: Arabidopsis CYCLIC NUCLEOTIDE GATED CHANNEL2 and Innate Immunity. The Plant Cell 19, 1081–1095.

Alqurashi M, Gehring C, Marondedze C. 2016. Changes in the Arabidopsis thaliana proteome implicate cAMP in biotic and abiotic stress responses and changes in energy metabolism. International Journal of Molecular Sciences 17.

Al-Younis I, Wong A, Gehring C. 2015. The Arabidopsis thaliana K+-uptake permease 7 (AtKUP7) contains a functional cytosolic adenylate cyclase catalytic centre. FEBS Letters 589, 3848–3852.

Al-Younis I, Wong A, Lemtiri-Chlieh F, Schmöckel S, Tester M, Gehring C, Donaldson L. 2018. The arabidopsis thaliana K+-uptake permease 5 (AtKUP5) contains a functional cytosolic adenylate cyclase essential for K+ transport. Frontiers in Plant Science 871, 1–15.

Ashton AR, Polya GM. 1978. Cyclic Adenosine 3′:5′-Monophosphate in Axenic Rye Grass Endosperm Cell Cultures 1. Plant Physiology 61, 718–722.

Azarashvili T, Krestinina O, Galvita A, Grachev D, Baburina Y, Stricker R, Evtodienko Y, Reiser G. 2009. Ca2+-dependent permeability transition regulation in rat brain mitochondria by 2′,3′-cyclic nucleotides and 2′,3′-cyclic nucleotide 3′-phosphodiesterase. American Journal of Physiology - Cell Physiology 296, 1428–1439.

Bhatt M, Di Iacovo A, Romanazzi T, Roseti C, Cinquetti R, Bossi E. 2022. The “www” of Xenopus laevis Oocytes: The Why, When, What of Xenopus laevis Oocytes in Membrane Transporters Research. Membranes 12.

Bianchet C, Wong A, Quaglia M, Alqurashi M, Gehring C, Ntoukakis V, Pasqualini S. 2019. An Arabidopsis thaliana leucine-rich repeat protein harbors an adenylyl cyclase catalytic center and affects responses to pathogens. Journal of Plant Physiology 232, 12–22.

Blanco E, Fortunato S, Viggiano L, de Pinto MC. 2020. Cyclic amp: A polyhedral signalling molecule in plants. International Journal of Molecular Sciences 21, 1–2.

Brown EG, Al-Najafi T, Newton RP. 1977. Cyclic nucleotide phosphodiesterase activity in Phaseolus vulgaris. Phytochemistry 16, 1333–1337.

Chatukuta P, Dikobe TB, Kawadza DT, Sehlabane KS, Takundwa MM, Wong A, Gehring C, Ruzvidzo O. 2018. An arabidopsis clathrin assembly protein with a predicted role in plant defense can function as an adenylate cyclase. Biomolecules 8, 1–15.

Chodasiewicz M, Kerber O, Gorka M, Moreno JC, Maruri-Lopez I, Minen RI, Sampathkumar A, Nelson ADL, Skirycz A. 2022. 2′,3′-cAMP treatment mimics the stress molecular response in Arabidopsis thaliana. Plant Physiology 188, 1966–1978.

Van Damme T, Blancquaert D, Couturon P, Van Der Straeten D, Sandra P, Lynen F. 2014. Wounding stress causes rapid increase in concentration of the naturally occurring 2′,3′-isomers of cyclic guanosine-and cyclic adenosine monophosphate (cGMP and cAMP) in plant tissues. Phytochemistry 103, 59–66.

Demidchik V, Cuin TA, Svistunenko D, Smith SJ, Miller AJ, Shabala S, Sokolik A, Yurin V. 2010. Arabidopsis root K+-efflux conductance activated by hydroxyl radicals: Single-channel properties, genetic basis and involvement in stress-induced cell death. Journal of Cell Science 123, 1468–1479.

Demidchik V, Shabala SN, Coutts KB, Tester MA, Davies JM. 2003. Free oxygen radicals regulate plasma membrane Ca2+-and K+-permeable channels in plant root cells. Journal of Cell Science 116, 81–88.

Diffley PE, Geisbrecht A, Newton RP, et al. 2001. Variation in isomeric products of a phosphodiesterase from the chloroplasts of Phaseolus vulgaris in response to cations. Plant Biosystems - An International Journal Dealing with all Aspects of Plant Biology 135, 143–156.

Ge SX, Jung D, Jung D, Yao R. 2020. ShinyGO: A graphical gene-set enrichment tool for animals and plants. Bioinformatics 36, 2628–2629.

Gehring C, Turek IS. 2017. Cyclic nucleotide monophosphates and their cyclases in plant signaling. Frontiers in Plant Science 8, 1–15.

Grau J, Franco-Zorrilla JM. 2022. TDTHub, a web server tool for the analysis of transcription factor binding sites in plants. Plant Journal 111, 1203–1215.

Jackson EK, Ren J, Mi Z. 2009. Extracellular 2′,3′-cAMP is a source of adenosine. Journal of Biological Chemistry 284, 33097–33106.

Jiang J, Fan LW, Wu WH. 2005. Evidences for involvement of endogenous cAMP in Arabidopsis defense responses to Verticillium toxins. Cell Research 15, 585–592.

Jin XC, Wu WH. 1999. Involvement of cyclic AMP in ABA-and Ca2+-mediated signal transduction of stomatal regulation in Vicia faba. Plant and Cell Physiology 40, 1127–1133.

Kosmacz M, Luzarowski M, Kerber O, et al. 2018. Interaction of 2’,3’-cAMP with Rbp47b Plays a Role in Stress Granule Formation. Plant physiology 177, 411–421.

Kozuleva MA, Petrova AA, Mamedov MD, Semenov AY, Ivanov BN. 2014. O2 reduction by photosystem I involves phylloquinone under steady-state illumination. FEBS Letters 588, 4364–4368.

Kurosaki F, Nishi A. 1993. Stimulation of Calcium Influx and Calcium Cascade by Cyclic AMP in Cultured Carrot Cells. Archives of Biochemistry and Biophysics 302, 144–151.

Leng Q, Mercier RW, Hua B-G, Fromm H, Berkowitz GA. 2002. Electrophysiological Analysis of Cloned Cyclic Nucleotide-Gated Ion Channels. Plant Physiology 128, 400–410.

Leng Q, Mercier RW, Yao W, Berkowitz GA. 1999. Cloning and first functional characterization of a plant cyclic nucleotide-gated cation channel. Plant Physiology 121, 753–761.

Li W, Luan S, Schreiber SL, Assmann SM. 1994. Cyclic AMP stimulates K+ channel activity in mesophyll cells of Vicia faba L. Plant Physiology 106, 957–961.

Lin PPC, Varner JE. 1972. Cyclic nucleotide phosphodiesterase in pea seedlings. Biochimica et Biophysica Acta (BBA) - Enzymology 276, 454–474.

Luo W, Xu Y, Cao J, et al. 2024. COLD6-OSM1 module senses chilling for cold tolerance via 2′,3′-cAMP signaling in rice. Molecular Cell 84, 4224–4238.e9.

Lüthje S, Möller B, Perrineau FC, Wöltje K. 2013. Plasma membrane electron pathways and oxidative stress. Antioxidants and Redox Signaling 18, 2163–2183.

Maathuis FJM, Sanders D. 2001. Sodium uptake in Arabidopsis roots is regulated by cyclic nucleotides. Plant Physiology 127, 1617–1625.

Moutinho A, Hussey PJ, Trewavas AJ, Malhó R. 2001. cAMP acts as a second messenger in pollen tube growth and reorientation. Proceedings of the National Academy of Sciences of the United States of America 98, 10481–10486.

Nikonorova N, Van Den Broeck L, Zhu S, Van De Cotte B, Dubois M, Gevaert K, Inzé D, De Smet I. 2018. Early mannitol-triggered changes in the Arabidopsis leaf (phospho)proteome reveal growth regulators. Journal of Experimental Botany 69, 4591–4607.

Ordoñez NM, Marondedze C, Thomas L, Pasqualini S, Shabala L, Shabala S, Gehring C. 2014. Cyclic mononucleotides modulate potassium and calcium flux responses to H2O2 in Arabidopsis roots. FEBS Letters 588, 1008–1015.

Ordoñez NM, Shabala L, Gehring C, Shabala S. 2013. Noninvasive microelectrode ion flux estimation technique (MIFE) for the study of the regulation of root membrane transport by cyclic nucleotides. Methods in Molecular Biology 1016, 95–106.

Paradiso A, Domingo G, Blanco E, Buscaglia A, Fortunato S, Marsoni M, Scarcia P, Caretto S, Vannini C, de Pinto MC. 2020. Cyclic AMP mediates heat stress response by the control of redox homeostasis and ubiquitin-proteasome system. Plant Cell and Environment 43, 2727–2742.

Pietrowska-Borek M, Nuc K. 2013. Both cyclic-AMP and cyclic-GMP can act as regulators of the phenylpropanoid pathway in Arabidopsis thaliana seedlings. Plant Physiology and Biochemistry 70, 142–149.

Ruzvidzo O, Gehring C, Wong A. 2019. New Perspectives on Plant Adenylyl Cyclases. Frontiers in Molecular Biosciences 6, 1–8.

Sabetta W, Vandelle E, Locato V, et al. 2019. Genetic buffering of cyclic AMP in Arabidopsis thaliana compromises the plant immune response triggered by an avirulent strain of Pseudomonas syringae pv. tomato. Plant Journal 98, 590–606.

Sampaio M, Neves J, Cardoso T, Pissarra J, Pereira S, Pereira C. 2022. Coping with Abiotic Stress in Plants—An Endomembrane Trafficking Perspective.

Shabala S, Pottosin I. 2014. Regulation of potassium transport in plants under hostile conditions: Implications for abiotic and biotic stress tolerance. Physiologia Plantarum 151, 257–279.

Tezuka T, Akita I, Yoshino N, Suzuki Y. 2007. Regulation of self-incompatibility by acetylcholine and cAMP in Lilium longiflorum. Journal of Plant Physiology 164, 878–885.

Thompson JE, Venegas FD, Raines RT. 1994. Energetics of Catalysis by Ribonucleases: Fate of the 2′,3′-Cyclic Phosphodiester Intermediate. Biochemistry 33, 7408–7414.

Tyc K, Kellenberger C, Filipowicz W. 1987. Purification and characterization of wheat germ 2’,3’-cyclic nucleotide 3’-phosphodiesterase. Journal of Biological Chemistry 262, 12994–13000.

Vannini C, Domingo G, Fiorilli V, Seco DG, Novero M, Marsoni M, Wisniewski-Dye F, Bracale M, Moulin L, Bonfante P. 2021. Proteomic analysis reveals how pairing of a Mycorrhizal fungus with plant growth-promoting bacteria modulates growth and defense in wheat. Plant Cell and Environment 44, 1946–1960.

Wang X, Xu M, Gao C, Zeng Y, Cui Y, Shen W, Jiang L. 2020. The roles of endomembrane trafficking in plant abiotic stress responses. Journal of Integrative Plant Biology 62, 55–69.

Wiśniewski JR. 2019. Filter Aided Sample Preparation – A tutorial. Analytica Chimica Acta 1090, 23–30.

Yu D, Song W, Tan EYJ, et al. 2022. TIR domains of plant immune receptors are 2′,3′-cAMP/cGMP synthetases mediating cell death. Cell 185, 2370–2386.e18.

Zelman AK, Dawe A, Gehring C, Berkowitz GA. 2012. Evolutionary and structural perspectives of plant cyclic nucleotide-gated cation channels. Frontiers in Plant Science 3, 1–13.

Zhao J, Guo Y, Fujita K, Sakai K. 2004. Involvement of cAMP signaling in elicitor-induced phytoalexin accumulation in Cupressus lusitanica cell cultures. New Phytologist 161, 723–733.

Zhao Y, Liu Y, Ji X, Sun J, Lv S, Yang H, Zhao X, Hu X. 2021. Physiological and proteomic analyses reveal cAMP-regulated key factors in maize root tolerance to heat stress. Food and Energy Security 10, 1–24.

Zhou L, Lan W, Jiang Y, Fang W, Luan S. 2014. A calcium-dependent protein Kinase interacts with and activates a calcium channel to regulate pollen tube growth. Molecular Plant 7, 369–376.

