## Supplemental file 1 for "Delineating plant responses to the 3’,5’- and 2’,3’-cAMP isomers"

**Supplementary protocol**

In order to overcome the problem of missing values without complex statistical analysis, we applied a hybrid approach as described by Nikonorova et al. 2018: we filtered data based on number of valid value and we applied a data analysis that treats intensity-based and presence/absence of data separately. Briefly the original, complete dataset containing log2transformed intensities was split in three subsets (Figure S1). The first subset consisted of proteins that were quantified in ≥ 75 % of the biological replicates in both control and cAMP-treated samples. These proteins were thus detected in at least 3 out of 4 biological replicates. The remaining missing values are considered MAR, Missing at Random, values and were imputated using an in-house R script based on KNN algorithm. The second subset contained proteins that were quantified in > 25% and < 75% of the biological replicates of at least one treatment. This group was named “unreliable” as only half of the replicates could be quantified and, was therefore excluded from further analysis. The third subset included proteins that were present in one sample (in more than 75 % of the replicates) and absent or below the detection threshold in another sample (in more than 75 % of the replicates). This subset contained “unique” proteins that are quite often incorrectly ignored and excluded from final results. Subsets of unique proteins were extracted and used as such, without any subsequent statistical analysis.
