## Supplemental figure S1 for "Delineating plant responses to the 3’,5’- and 2’,3’-cAMP isomers"

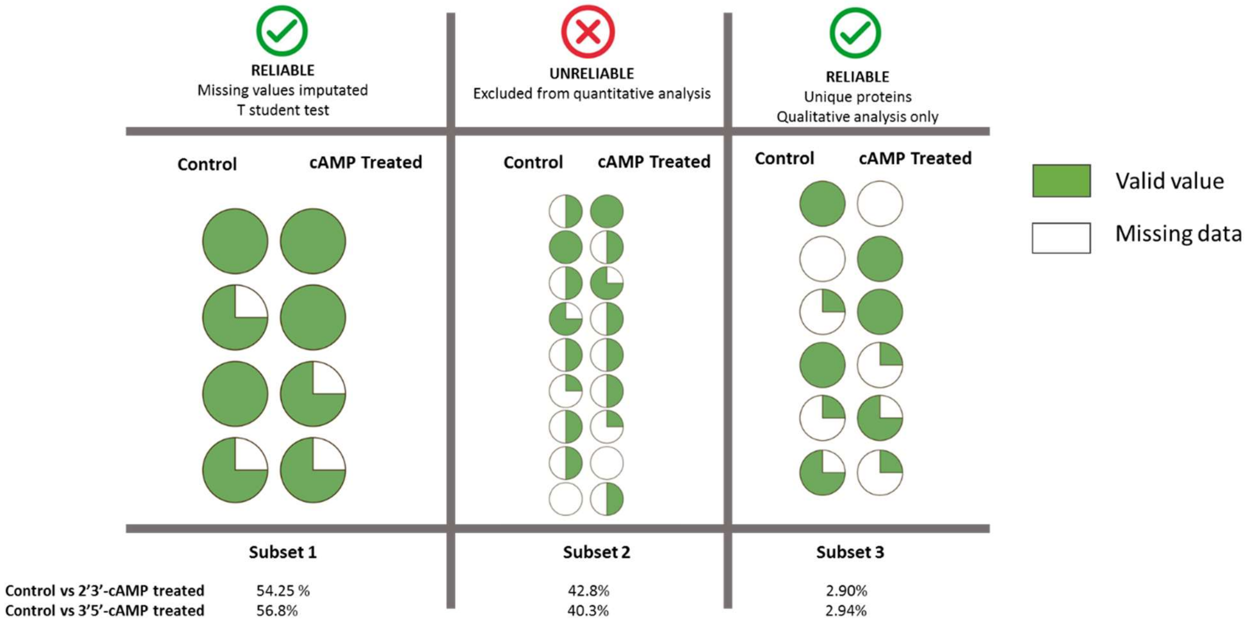


**Figure S1**. Visual explanation for the subsets described in the Materials and methods. The values detected for each protein are represented in a circle and each quarter of a circle represents a biological repeat. For example, a half green circle means that only two valid values were detected. Depending on the percentage of valid values each protein was subdivided in one of the three subsets. The percentage of each dataset and subset are indicated.
